# The Limbic Overload Hypothesis of Hypomanic Vulnerability: A Dynamic Biosocial Perspective

**DOI:** 10.64898/2026.05.25.727695

**Authors:** Helen Pushkarskaya, Godfrey Pearlson, Christopher Pittenger

## Abstract

Hypomanic tendencies are associated with elevated goal-directed behavior, creativity, charisma, sociability, and entrepreneurial drive, but also with mood instability, irritability, impulsive persistence, and elevated risk for bipolar disorder and other psychopathology. Existing models often emphasize unidimensional constructs such as reward sensitivity or behavioral activation, yet these approaches incompletely capture the dynamic and often contradictory nature of the hypomanic temperament. We propose the Limbic Overload Hypothesis of Hypomanic Vulnerability, a dynamic biosocial framework suggesting that hypomanic tendencies reflect a persistent pattern of elevated engagement despite potential loss, coupled with reduced integration of negative emotional experience into subsequent behavioral regulation. Over time, this pattern may contribute to progressive “limbic overload,” characterized by increasing emotional dysregulation, hypersensitivity to salient experiences, and vulnerability to psychopathology. Integrating evidence from personality research, affective neuroscience, and preliminary neuroimaging findings, we propose a dynamic cortico-limbic model linking prefrontal-limbic coordination, loss tolerance, emotional updating, and social reinforcement cycles. Preliminary pilot data suggest that individual differences in hypomanic tendencies are reflected not simply in baseline cortico-limbic organization, but in dynamic neural reconfiguration across pre-task resting-state → task → post-task resting-state transitions during loss-related decision making. Specifically, elevated hypomanic tendencies were associated with persistently elevated tolerance of potential losses and reduced integration of negative emotional information into subsequent behavioral regulation. We further propose that social connectedness and cognitive-emotional integration may mitigate progressive limbic overload and contribute to resilience. Together, this framework generates experimentally testable predictions regarding the neural, behavioral, and social processes underlying hypomanic vulnerability and resilience.

## Introduction

No two people respond to challenge, reward, loss, and social feedback in the same way. Some individuals persist through uncertainty and repeated setbacks with unusual energy and engagement, whereas others become overwhelmed or withdraw. Understanding how such persistent individual differences emerge requires models that account not only for static traits, but also for dynamic interactions among neural regulation, behavior, and social experience over time. Hypomanic tendencies provide a compelling model system for investigating these processes.

Individuals with elevated hypomanic tendencies exhibit a striking combination of characteristics that appear psychologically difficult to reconcile within a single coherent temperament profile: high energy, charisma, creativity, sociability, and goal orientation, but also irritability, impulsive persistence, inflated self-confidence, interpersonal instability, and relative disregard for social norms. Although it is not immediately obvious why these tendencies should co-occur, empirical studies consistently demonstrate that they cluster together within the hypomanic temperament (Schalet et al., 2011). Classical clinical descriptions of elevated MMPI Hypomania (Ma) scores emphasized unstable activation patterns, fluctuating emotional intensity, exaggerated self-confidence, impulsive engagement, and socially disinhibited behavior (McKinley & Hathaway, 1944). Similarly, assessment interviews conducted by Eckblad and Chapman (1986) described periods of unusually elevated activity alternating with relative passivity, affective states characterized by simultaneous positive and irritable mood, and contradictory interpersonal patterns later described as “gregarious social misfits” (Pushkarskaya et al., 2021). To operationalize these tendencies dimensionally in the general population, Eckblad and Chapman developed the Hypomanic Personality Scale (HPS), which remains one of the most widely used measures of hypomanic temperament. Factor analytic studies of the HPS consistently identify three intercorrelated dimensions—social vitality, mood volatility, and excitement—suggesting that hypomanic temperament reflects a structured but internally heterogeneous configuration of partially opposing characteristics rather than a single elevated trait dimension (Schalet et al., 2011).

Elevated hypomanic tendencies represent a temporally stable cognitive-behavioral pattern that is distinct from a hypomanic episode. Approximately 50–70% of individuals with elevated hypomanic tendencies experience a hypomanic episode during their lifetime (Eckblad & Chapman, 1986; Kwapil et al., 2000), and roughly 15% develop bipolar-spectrum illness, compared to approximately 2.4% of the general population (Faedda et al., 2019; Homish et al., 2013). Elevated hypomanic tendencies are also associated with increased risk for mood disorders, substance abuse, and other forms of psychopathology (Eckblad & Chapman, 1986; Kwapil et al., 2000; Meyer & Hautzinger, 2003). At the same time, many individuals with elevated hypomanic tendencies do not develop severe mental illness and instead demonstrate high levels of adaptive functioning. Hypomanic tendencies have long been linked to creativity, elevated motivation, and increased goal-directed behavior (Clare, 1997). In our prior work across three independent populations (total N = 2,200), elevated hypomanic tendencies were associated with a history of business creation, particularly among serial entrepreneurs (Pushkarskaya et al., 2021). Importantly, however, this association was observed primarily among individuals with relatively preserved social adjustment, indexed by lower MMPI Psychopathic Deviate scores (Willerman et al., 1992). Together with prior longitudinal findings (Kwapil et al., 2000), these observations suggest that the same underlying temperament profile may contribute to markedly different long-term outcomes depending on how emotional, interpersonal, and regulatory processes evolve over time. Developing interventions that reduce risk while preserving adaptive functioning therefore requires a clearer mechanistic understanding of the processes that contribute both to vulnerability and to resilience within the hypomanic temperament.

Many contemporary models attempt to explain hypomanic tendencies using relatively unidimensional constructs such as behavioral activation, reward sensitivity, impulsivity, or sensation seeking (Baas et al., 2020; de Oliveira et al., 2019; Johnson, 2005; Johnson et al., 2012). These approaches align with dimensional psychiatry frameworks emphasizing decomposition of psychopathology into simpler transdiagnostic processes. However, several observations suggest that such models may incompletely capture the structure and dynamics of the hypomanic temperament. Instability itself has long been recognized as a defining feature of hypomanic personality and bipolar-spectrum vulnerability. Classical descriptions of elevated MMPI Hypomania (Ma) scores emphasized persistently fluctuating activation, emotional variability, irritability, impulsive engagement, and unstable interpersonal functioning rather than a unitary elevation in reward sensitivity or positive affect alone (McKinley & Hathaway, 1944). Similarly, factor analytic studies of the Hypomanic Personality Scale (HPS) consistently support a multidimensional structure composed of partially distinct but intercorrelated factors, including social vitality, mood volatility, and excitement (Schalet et al., 2011). More broadly, the central phenomenon requiring explanation may not simply be the presence of individual hypomanic characteristics, but why seemingly contradictory emotional, behavioral, and interpersonal tendencies persistently co-occur within a chronically unstable temperament profile. Understanding hypomanic vulnerability may therefore require shifting the central question from which isolated traits are associated with hypomanic tendencies to what mechanisms both give rise to this persistent co-occurrence and sustain the enduring instability that defines the hypomanic temperament.

To investigate these questions, we present convergent evidence from two complementary studies examining hypomanic tendencies across behavioral and neural levels of analysis. First, using longitudinal self-report data, we examine whether components of hypomanic temperament exhibit stable temporal coupling over time. Second, using functional neuroimaging data, we examine whether elevated hypomanic tendencies are associated with variations in neural dynamics within brain networks implicated in emotional regulation, salience processing, and goal-directed behavior. Together, these findings are used to develop a dynamic biosocial model of hypomanic vulnerability aimed at explaining why seemingly contradictory characteristics persistently co-occur within the hypomanic temperament, how these interacting processes may under certain conditions contribute to emotional and limbic dysregulation, and what regulatory mechanisms may promote resilience and adaptive functioning.

## Methods

### Study 1. Longitudinal Organization of Hypomanic Temperament

#### Participants and Procedure

Participants were 165 undergraduate students (85 females; mean age = 18 years, SD = 1) recruited prior to entry into college. Data were collected across two waves of online assessment designed capture a major developmental and social transition period. Wave 1 assessment occurred during August 2018, approximately one week prior to the beginning of college, whereas Wave 2 assessment occurred during April 2020, during participants’ fourth academic semester following approximately two years of academic and social adaptation to university life. All participants provided informed consent in accordance with institutional review board procedures and were compensated for their time.

#### Measures

The Hypomanic Personality Scale (HPS; Eckblad & Chapman, 1986) is a self-report measure designed to assess stable hypomanic personality characteristics and vulnerability to bipolar-spectrum psychopathology in nonclinical populations. Prior factor analytic studies support a multidimensional structure composed of three intercorrelated dimensions: Social Vitality, Mood Volatility, and Excitement. The HPS has demonstrated strong psychometric properties and predictive validity for bipolar-spectrum outcomes and related psychopathology (Eckblad & Chapman, 1986; Kwapil et al., 2000).

The Sensitivity to Punishment and Sensitivity to Reward Questionnaire (SPSRQ; Torrubia et al., 2001) is a self-report measure based on Gray’s Reinforcement Sensitivity Theory designed to assess individual differences in behavioral inhibition and behavioral activation tendencies. The SPSRQ has demonstrated good reliability and validity across clinical and nonclinical populations (Caseras et al., 2003; Torrubia et al., 2001).

Depression Anxiety Stress Scale (DASS-21; Lovibond, 1995) is a 21-item self-report measure assessing symptoms of depression, anxiety, and stress over the preceding week. The DASS-21 yields both subscale scores and a total psychological distress score and has demonstrated strong reliability and convergent validity in both clinical and community samples (Antony et al., 1998; Lovibond, 1995).

Mood Disorder Questionnaire (MDQ; Hirschfeld et al., 2000) is a self-report screening measure designed to assess lifetime history of manic and hypomanic symptoms and associated functional impairment. The MDQ includes items assessing elevated mood, increased energy, decreased need for sleep, racing thoughts, distractibility, increased sociability, impulsive/risky behavior, and other symptoms characteristic of bipolar-spectrum vulnerability (Hirschfeld et al., 2000).

Student Adaptation to College Questionnaire (SACQ; Baker & Siryk, 1984) is a self-report measure designed to assess academic, emotional, social, and institutional adjustment to university life. The SACQ includes subscales assessing social adjustment and emotional adjustment, which were used in the present analyses. The SACQ has demonstrated good psychometric properties and predictive validity for college adjustment and psychosocial functioning (Baker & Siryk, 1984).

Resilience Scale for Adults (RSA; Friborg et al., 2003) is a self-report measure designed to assess protective factors associated with psychological resilience and adaptive functioning. The RSA assesses dimensions including perception of self, planned future, social competence, family cohesion, social resources, and structured style. The RSA has demonstrated good reliability and construct validity across clinical and nonclinical populations (Friborg et al., 2003).

Demographic characteristics assessed at Wave 1 included age, sex, parental income, and race. Hypomanic tendencies were assessed at both Wave 1 and Wave 2. Additionally, at Wave 1 participants completed measures assessing reward sensitivity, punishment sensitivity, and demographic characteristics, whereas at Wave 2 participants completed measures assessing psychological distress, adjustment to student life, and resilience strategies.

#### Longitudinal Structural Equation Modeling

Study 1 examined longitudinal relationships among dimensions of hypomanic temperament, reward sensitivity, social-regulatory processes, and later psychological outcomes using structural equation modeling (SEM). Analyses focused on the three established HPS dimensions—Social Vitality, Mood Volatility, and Excitement—modeled using observed subscale scores derived from prior factor analytic work (Schalet et al., 2011).

Two complementary longitudinal models were estimated. The first examined relationships among reward sensitivity, hypomanic temperament dimensions, and later psychological distress outcomes, including depression, anxiety, and stress symptoms assessed by the DASS-21 and mood dysregulation assessed by MDQ. The second examined relationships among baseline hypomanic temperament dimensions, resilience-related processes, emotional and social adjustment, and corresponding Wave 2 hypomanic temperament dimensions.

Structural equation models were estimated using maximum likelihood estimation with bootstrapping (1,000 resamples). Model fit was evaluated using chi-square statistics, root mean square error of approximation (RMSEA), standardized root mean square residual (SRMR), and P-close. Standardized path coefficients are reported throughout.

### Study 2. Neural Dynamics Associated with Hypomanic Tendencies

#### Participants

Nineteen healthy young adults were recruited from the local community and university population as part of a larger study examining emotional decision-making and neural dynamics associated with hypomanic tendencies. Participants were between 23 and 56 years of age (M = 38, SD = 8; 10 females). Exclusion criteria included a history of neurological illness, major psychiatric disorder, substance dependence, and standard MRI contraindications. All participants provided written informed consent in accordance with institutional review board procedures and were compensated for their participation.

Prior to neuroimaging assessment, participants completed self-report measures assessing hypomanic tendencies, behavioral activation and inhibition, sensation seeking, and demographic characteristics.

#### Measures

Hypomanic Personality Scale (HPS; Eckblad & Chapman, 1986) is a self-report measure designed to assess stable hypomanic personality characteristics and vulnerability to bipolar-spectrum psychopathology in nonclinical populations. The HPS assesses traits including elevated energy, social vitality, impulsive excitement, mood volatility, and increased goal-directed behavior. Prior work has demonstrated strong psychometric properties and predictive validity for bipolar-spectrum outcomes and related psychopathology (Eckblad & Chapman, 1986; Kwapil et al., 2000).

Behavioral Inhibition System/Behavioral Activation System scales (BIS/BAS; Carver & White, 1994) is a self-report measure based on Gray’s Reinforcement Sensitivity Theory. The BIS scale assesses sensitivity to punishment and behavioral inhibition, whereas the BAS scales assess reward responsiveness, drive, and reward-motivated behavior. The BIS/BAS scales have demonstrated good reliability and validity across clinical and nonclinical populations (Carver & White, 1994).

Sensation Seeking Scale (SSS; Zuckerman, 1994) is a self-report measure designed to assess preference for novel, intense, and stimulating experiences, including willingness to engage in risk-taking behaviors despite potential negative consequences. The SSS has demonstrated good psychometric properties and has been widely used in studies of impulsivity, reward processing, and affective psychopathology (Zuckerman, 1994, 2007).

Demographic characteristics included age, sex, race, and ethnicity.

#### Experimental Task

Participants completed a modified Risk and Ambiguity (R&A; *Fig. 1*) decision-making task previously described by Levy et al. (2010). On each trial, participants chose between a certain monetary option (±$10) and a lottery option varying in both outcome magnitude and outcome probability. Potential gains and losses ranged from $10 to $100. In risky trials, outcome probabilities were explicitly specified (13%, 25%, 38%, 50%, or 75%), whereas in ambiguous trials probability information was partially occluded to introduce uncertainty regarding outcome likelihood.

**Figure 1.**
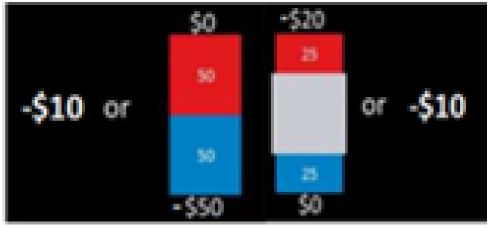
Sample stimuli for the Risk and Ambiguity task.

Trials were organized into separate gain and loss blocks, allowing examination of decision-making behavior under positive and negative outcome contexts. The task was designed to assess persistence of goal-directed engagement despite potential losses and uncertainty, processes hypothesized to be particularly relevant to hypomanic vulnerability. Behavioral indices focused on individual differences in tolerance for potential losses, operationalized as the proportion of loss trials in which participants elected to gamble rather than select the certain option.

Two resting-state scans were acquired, one prior to task performance (Rest I) and one immediately following the task (Rest II), allowing examination of how intrinsic neural connectivity patterns both influenced and were altered by task-related emotional and decision-making processes. At the end, one trial is played for real money. Winnings or losses are added to initial endowment of $100.

#### MRI Acquisition

Imaging data were acquired on a Siemens 3T MRI scanner at the Yale Magnetic Resonance Research Center using Human Connectome Project (HCP)-style acquisition protocols (Glasser, Smith, et al., 2016; Glasser et al., 2013; Harms et al., 2018; Yang et al., 2014). High-resolution T1-weighted and T2-weighted anatomical images were collected for anatomical localization and spatial normalization. Functional blood oxygenation level-dependent (BOLD) images were acquired using a multiband accelerated gradient-echo echo-planar imaging (EPI) sequence (TR = 720 ms, TE = 36 ms, voxel size = 2.1 mm isotropic, 64 axial slices, multiband factor = 8).

Two resting-state scans were acquired, one prior to task performance (Rest I) and one following completion of the task (Rest II), each consisting of 420 frames (∼ 5 min). Task-based fMRI data were acquired during performance of the Risk and Ambiguity decision-making task.

#### fMRI Preprocessing

Imaging data were preprocessed using the Human Connectome Project (HCP) minimal preprocessing pipeline (Glasser et al., 2013) implemented through the Quantitative Neuroimaging Environment & Toolbox (QuNex; (Ji et al., 2023)). Preprocessing included motion correction, spatial normalization, co-registration of functional and anatomical images, and artifact removal. Motion-related artifacts were additionally addressed using framewise displacement and signal-intensity–based scrubbing procedures consistent with prior recommendations for resting-state fMRI analyses (Power et al., 2013). Frames exceeding motion or signal-intensity thresholds, along with adjacent frames, were excluded from analyses (Anticevic et al., 2012).

A trained research assistant visually inspected all scans for co-registration quality, excessive movement, signal loss, magnetic field inhomogeneities, and potential anatomical abnormalities. One female participant was removed due to excessive motion. A temporal high-pass filter (0.008 Hz) was applied to remove low-frequency scanner drift. Nuisance regression additionally removed signals derived from ventricles, deep white matter, global gray matter signal, motion parameters, and their first derivatives from gray matter BOLD time series.

#### Functional connectivity measures

Neuroimaging analyses focused on regions implicated in emotional reactivity, emotional memory integration, and emotion regulation, including the amygdala (AMY), hippocampus (Hipp), and ventromedial prefrontal cortex (vmPFC) (Etkin et al., 2015). Analyses focused on the right hemisphere, which has been more strongly implicated in emotional and particularly negative emotional processing (Campbell, 1982; Mandal et al., 1991). Regions of interest (ROIs) were defined a priori using the Glasser parcellation (Glasser, Coalson, et al., 2016). The right amygdala was examined as a region associated with emotional salience and reactivity (Ochsner et al., 2009), the right hippocampus as a region implicated in emotional episodic memory processing (Phelps, 2004), and vmPFC as a region associated with reward valuation, emotion regulation, and cognitive-emotional integration (Critchley, 2005; Gu et al., 2013; Hiser & Koenigs, 2018; Winecoff et al., 2013).

Resting-state functional connectivity analyses were conducted separately for resting-state scans acquired before (Rest I) and after (Rest II) completion of the Risk and Ambiguity task. To compute resting-state functional connectivity measures, mean BOLD time series were first extracted and averaged across all voxels within each ROI. Pairwise Pearson correlations were then computed between ROI time series for Rest I and Rest II separately. The relations between intrinsic brain activity and operations underlying perception and behavior is bi-directional (Sadaghiani & Kleinschmidt, 2013): fluctuations in intrinsic activity reflect the past history of the system and influence present and future operations. Thus, our design allowed examination of intrinsic functional connectivity patterns both preceding task engagement and following exposure to emotionally and motivationally salient decision-making experiences.

#### Neural activation measures

Task-related neural activation was estimated using general linear models (GLMs) implemented separately for each participant. Gain and loss trial events were modeled as separate regressors and convolved with the canonical hemodynamic response function. Nuisance regressors derived from preprocessing procedures, including motion parameters and physiological noise estimates, were included as covariates in all models.

First-level contrast estimates corresponding to gain and loss conditions were computed for each participant and entered into group-level random-effects analyses. Mean activation estimates for gain and loss contrasts were then extracted from predefined anatomical regions of interest (ROIs), including the right amygdala (rAMY), right hippocampus (rHipp), and nucleus accumbens (NAcc). The amygdala and hippocampus were selected based on their roles in emotional reactivity and emotional memory integration, whereas the NAcc was included because of its established involvement in reward and loss valuation processes (Carter et al., 2009).

Activation estimates derived from loss-related contrasts were used in subsequent analyses examining relationships among neural responses to potential losses, behavioral loss tolerance, resting-state functional connectivity, and hypomanic tendencies.

#### Longitudinal Structural Equation Modeling

Structural equation modeling (SEM) was used to examine relationships among task-related changes in resting-state functional connectivity, task-related neural activation during loss processing, behavioral loss tolerance, and hypomanic tendencies. The model focused on rest–task–rest dynamics among the ventromedial prefrontal cortex (vmPFC), right amygdala (rAMY), and right hippocampus (rHipp), as well as task-related activation within the rAMY, rHipp, and nucleus accumbens (NAcc) during loss trials.

Primary models examined whether intrinsic functional connectivity patterns during pre-task rest (Rest I) predicted neural activation within the rAMY, rHipp, and NAcc during loss trials, and whether neural activation during loss processing subsequently predicted changes in functional connectivity observed from pre-task to post-task rest (Rest II). Functional connectivity change scores were computed as Rest II minus Rest I for each ROI pair. This modeling framework was designed to examine relationships among intrinsic neural organization, emotional responses to potential losses, behavioral persistence under loss, and post-task cortico-limbic regulatory states.

Additional models examined associations between resting-state functional connectivity measures, behavioral loss tolerance, hypomanic tendencies, behavioral activation, and sensation-seeking traits. Structural equation models were estimated using maximum likelihood estimation with bootstrapping (1,000 resamples). Model fit was evaluated using chi-square statistics, root mean square error of approximation (RMSEA), standardized root mean square residual (SRMR), and P-close. Standardized path coefficients are reported throughout.

## Results

### Study 1. Longitudinal Organization of Hypomanic Temperament

#### Reward Sensitivity, Hypomanic Tendencies, and Future Psychological Distress

The longitudinal SEM examining relationships among reward sensitivity, hypomanic tendencies, and later psychological distress demonstrated good model fit (CFI = 0.99, CMIN/DF = 1.70, RMSEA = 0.076, PCLOSE = 0.27; *Fig. 2*). Greater sensitivity to rewards at Wave 1 was positively associated with hypomanic tendencies (β = 0.29, p < 0.001), whereas sensitivity to punishment was negatively associated with hypomanic tendencies (β = −0.15, p < 0.05).

**Figure 2.**
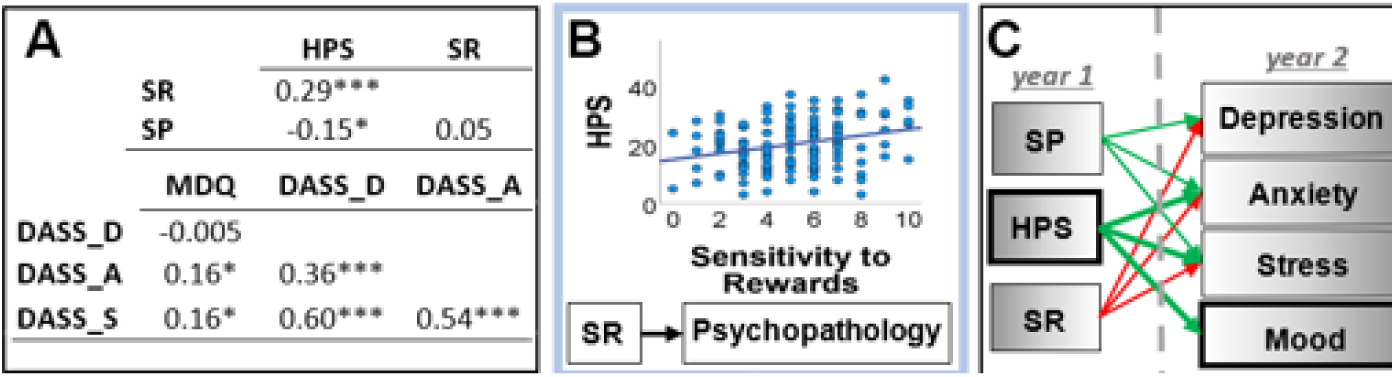
Reward Sensitivity, Hypomanic Tendencies, and Future Distress.

Critically, hypomanic tendencies predicted greater anxiety (β = 0.36, p < 0.001), stress (β = 0.60, p < 0.001), and mood-related symptoms (β = 0.54, p < 0.001) at Wave 2 even after accounting for individual differences in reward and punishment sensitivity. Direct associations between reward sensitivity and later psychological distress were comparatively weak and inconsistent across outcomes.

#### Longitudinal trajectories of hypomanic tendencies

The longitudinal SEM examining relationships among hypomanic temperament dimensions, adjustment processes, and resilience-related factors demonstrated acceptable model fit (CFI = 0.97, CMIN/DF = 1.82, RMSEA = 0.11, PCLOSE = 0.054; *Fig. 3*). The three HPS dimensions—Social Vitality, Mood Volatility, and Excitement—remained positively correlated within both waves, indicating persistent coupling among partially distinct hypomanic temperament components across time. Mood Volatility and Excitement demonstrated particularly strong associations at both Wave 1 (r = 0.50, p < 0.001) and Wave 2 (r = 0.42, p < 0.01), despite exhibiting opposite relationships with social and emotional adjustment outcomes. Emotional and social adjustment were themselves positively correlated, yet Mood Volatility negatively predicted both emotional and social adjustment, whereas Excitement positively predicted social adjustment. This persistent co-occurrence of partially opposing processes is consistent with classical clinical descriptions of hypomanic temperament as characterized by enduring instability rather than isolated elevations in specific traits.

**Figure 3.**
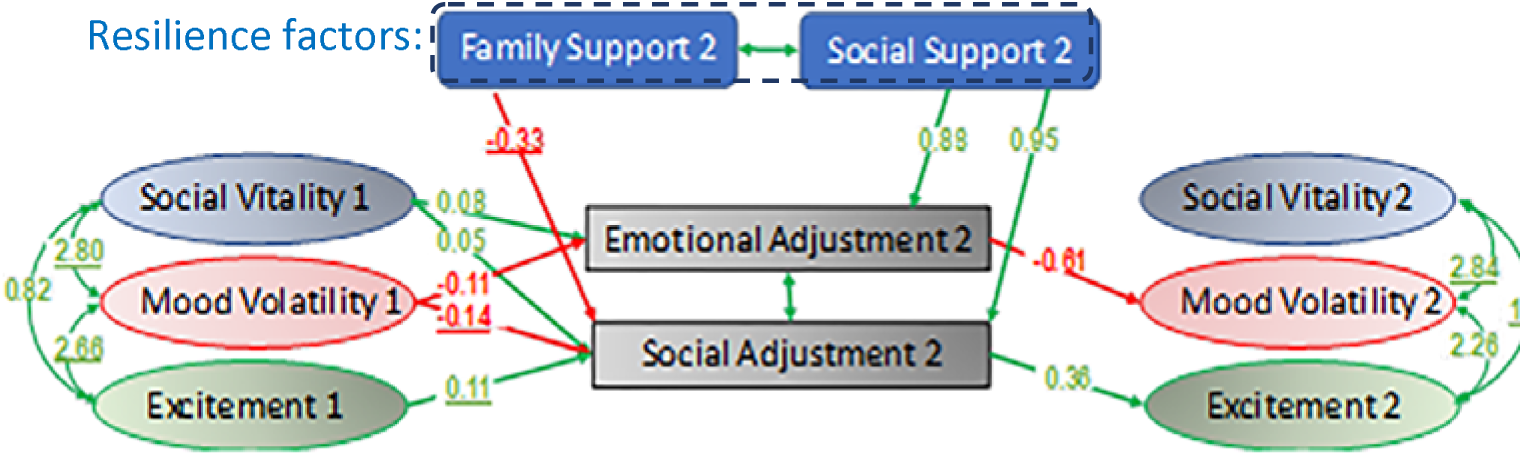
The corresponsive principle of development of hypomanic temperment. Emotional Adjustment and Social Adjustment are subscales of SACQ; Family Support and Social Support are subscales of the Resilience Scale. <u>Shown:</u> significant relations at p≤0.05: **positive** and **negative**. <u>Not</u> <u>shown:</u> Social Vitality cov. (t1,t2) = 7.7 p < 0.001; Mood Vitality cov. (t1,t2) = 1.2 p =0.21; Excitement cov. (t1,t2)= 0.9 p = 0.02; CFI = 0.97, CMIN/DF = 1.82 with N = 75, RMSEA = 0.11. PCLOSE =

Longitudinal analyses additionally revealed distinct adaptive and maladaptive feedback patterns consistent with the corresponsive principle of personality development, which proposes that the traits that steer individuals toward particular life situations are themselves reinforced and amplified by those environments (Harms, 2019). Greater Mood Volatility at Wave 1 predicted poorer emotional adjustment at Wave 2, which in turn predicted greater Mood Volatility at Wave 2, consistent with a maladaptive emotional feedback cycle. In contrast, greater Excitement at Wave 1 predicted stronger social adjustment at Wave 2, which subsequently predicted greater Excitement at Wave 2, consistent with a positive social-engagement feedback cycle.

In contrast to Mood Volatility and Excitement, Social Vitality did not substantially participate in either longitudinal feedback cycle and primarily demonstrated stability across waves. Social support from peers was positively associated with emotional and social adjustment outcomes, whereas greater family-resource dependence demonstrated negative associations with social adjustment during young adults’ transition to independence.

Since Mood Volatility and Excitement strongly correlated both in the present data and in prior studies (Schalet et al., 2011), these adaptive and maladaptive developmental cycles may unfold concurrently within the hypomanic temperament, potentially contributing to the characteristic combination of elevated engagement, instability, and behavioral intensity often described clinically as the “hypomanic edge” (Gartner, 2008).

### Study 2. Neural Dynamics Associated with Hypomanic Tendencies

Consistent with prior findings linking ventral striatal responses to salience and loss sensitivity during decision-making under uncertainty (Levy et al., 2010; Tom et al., 2007), greater NAcc activation during loss trials was associated with reduced behavioral loss tolerance, such that individuals with stronger NAcc responses were less likely to gamble under potential losses.

Given the modest sample size of this pilot neuroimaging study, SEM analyses were intended primarily as a proof of concept for modeling hypomanic tendencies in the context of dynamic cortico-limbic reconfiguration across rest–task–rest transitions. The SEM demonstrated good model fit (χ²(22) = 12.69, p = 0.94, CFI = 1.0, CMIN/DF = 0.58, RMSEA = 0.00, PCLOSE = 0.95; *Fig. 4*). More broadly, the overall rest–task–rest framework is consistent with prior work demonstrating that intrinsic resting-state connectivity predicts subsequent task-related neural responses and emotional processing dynamics (Mennes et al., 2010; Sakaki et al., 2013).

**Figure 4.**
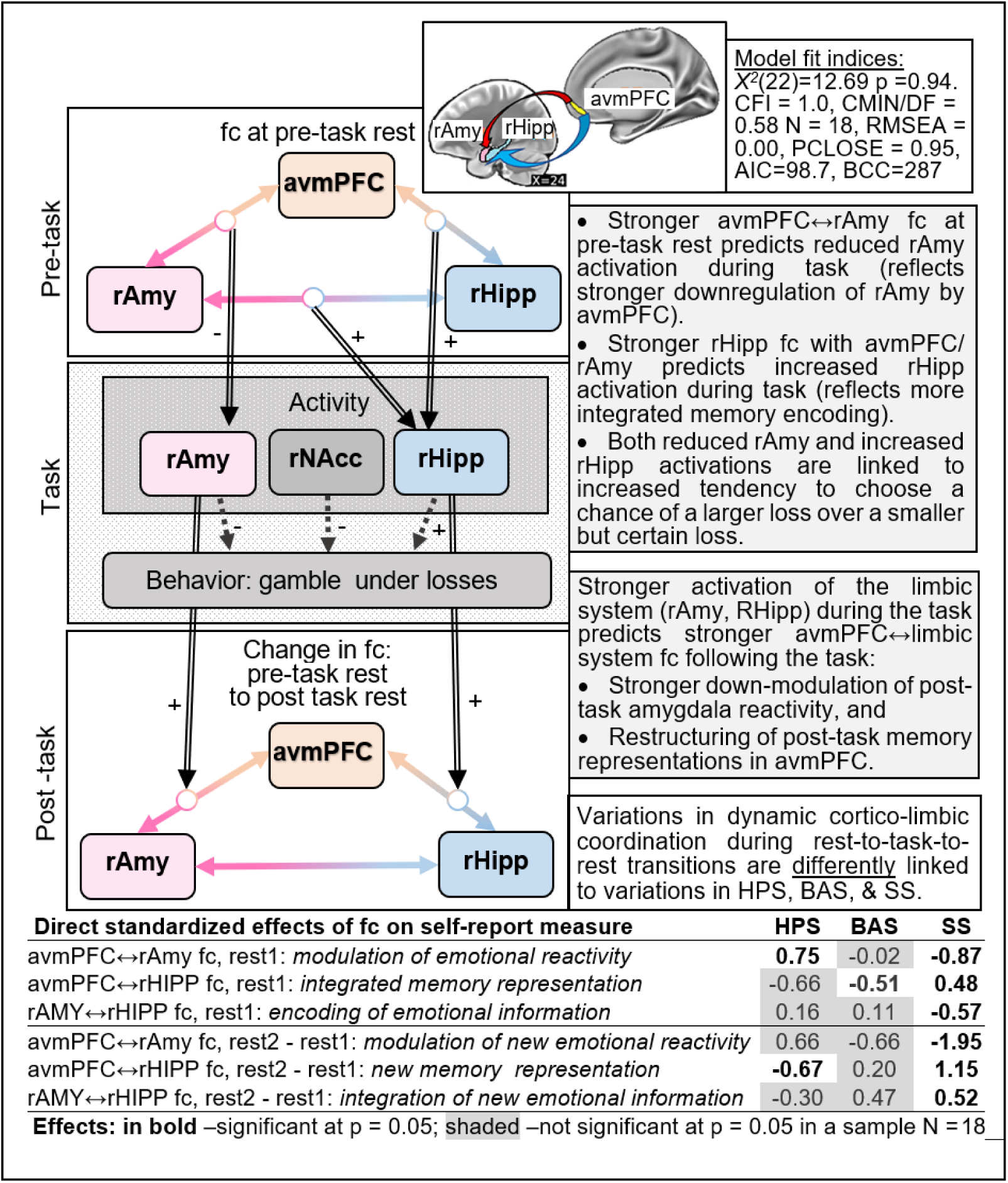
The dynamic Prefrontal-Limbic Coordination model.

Two partially parallel cortico-limbic dynamic patterns were observed. First, stronger pre-task avmPFC↔rAmy functional connectivity predicted reduced rAmy activation during loss processing, consistent with stronger baseline down-modulation of limbic emotional reactivity. Prior work has similarly linked stronger vmPFC↔amygdala coupling to reduced emotional reactivity and greater regulatory capacity (Banks et al., 2007; Eden et al., 2015; Motzkin et al., 2015). In our data, reduced rAmy activation during the task was additionally associated with greater willingness to gamble under potential losses. In turn, reduced rAmy activation predicted smaller increases in avmPFC↔rAmy connectivity during post-task rest, suggesting reduced post-task emotional regulatory reconfiguration following exposure to potential losses. Together, this pattern may reflect a persistent engagement-related process characterized by reduced emotional sensitivity to potential losses supporting continued goal-directed behavior.

Second, stronger baseline integration of rHipp with avmPFC and rAmy (i.e., high baseline functional connectivity) predicted greater rHipp activation during loss processing. Greater willingness to gamble under potential losses was associated with greater rHipp activation during the task, consistent with greater engagement of hippocampal systems involved in encoding and integration of emotionally salient experience. In contrast to the amygdala pathway, greater rHipp activation subsequently predicted stronger post-task avmPFC↔rHipp connectivity, consistent with greater post-task integration or restructuring of recent emotional experience following task exposure. This interpretation is broadly consistent with prior work linking hippocampal-amygdala connectivity to emotional memory integration and persistence of emotionally salient experiences (de Voogd et al., 2017; Dolcos et al., 2004; McGaugh, 2004).

Hypomanic tendencies were associated with distinct variations in cortico-limbic dynamics that were not fully captured by unidimensional measures of behavioral activation or sensation seeking. Higher HPS scores were associated with stronger baseline downmodulation of amygdala reactivity together with reduced post-task integration of hippocampal emotional experience within cortico-limbic regulatory circuits. In contrast, BAS scores were associated primarily with reduced baseline integration of the rHipp within the cortico-limbic network, whereas sensation-seeking was associated with broader dysregulation of vmPFC–rAmy–rHipp circuit dynamics (see *Fig. 4*).

## Discussion

Across two complementary studies, the present findings suggest that hypomanic tendencies may reflect a dynamic organizational process characterized by persistently elevated engagement despite increasing emotional dysregulation over time. Longitudinal analyses demonstrated stable coupling among partially opposing dimensions of hypomanic temperament together with parallel adaptive and maladaptive developmental trajectories. In parallel, pilot neuroimaging findings demonstrated that elevated hypomanic tendencies were associated with distinct patterns of cortico-limbic reconfiguration across rest→task→post-task transitions during loss-related decision making. Together, these findings suggest that the enduring instability characteristic of hypomanic temperament may emerge not simply from isolated elevations in reward sensitivity or impulsivity, but from dynamic interactions among persistent goal-directed engagement, emotional processing, and experience-based adaptation over time.

Existing models of hypomanic tendencies have primarily emphasized relatively unidimensional processes such as elevated reward sensitivity, behavioral activation, impulsivity, or sensation seeking (Baas et al., 2020; de Oliveira et al., 2019; Johnson, 2005; Johnson et al., 2012). These frameworks help explain persistent approach behavior, elevated goal pursuit, novelty seeking, and increased willingness to engage under conditions of uncertainty. However, they less readily explain several defining features of the hypomanic temperament, including the persistent co-occurrence of adaptive and maladaptive characteristics, marked emotional and interpersonal instability, and the observation that many individuals with elevated hypomanic tendencies simultaneously demonstrate elevated achievement and increased vulnerability to psychopathology (Eckblad & Chapman, 1986; Kwapil et al., 2000; Schalet et al., 2011). In the present studies, hypomanic tendencies were associated not simply with generalized reward-related traits, but with distinct longitudinal and neural dynamic patterns that were not fully captured by BAS or sensation-seeking measures alone. Specifically, longitudinal analyses demonstrated persistent coupling among partially opposing dimensions of hypomanic temperament together with parallel adaptive and maladaptive developmental trajectories, whereas pilot neuroimaging findings demonstrated stronger baseline downmodulation of amygdala reactivity together with reduced post-task integration of hippocampal emotional experience within cortico-limbic regulatory circuits during loss-related decision making. Together, these findings suggest that understanding hypomanic vulnerability may require moving beyond static trait elevations toward models focused on dynamic interactions among persistent engagement, emotional updating, and regulatory adaptation over time.

Based on these models, we propose that stably elevated loss tolerance may help explain multiple aspects of elevated hypomanic tendencies (*Fig. 5*). Stably elevated loss tolerance makes these individuals quick to engage in goal-directed activities, even in the presence of potential loss. Behaviorally, this pattern may manifest as unusual intensity and persistence in goal-directed engagement. They do not slow down to process after exposure to potential loss, potentially leading to persistent social and goal-oriented engagement (a contributing factor to the virtuous cycle depicted in *Fig. 3&5*). This decreased integration of negative experience (encoded in Hipp) into the value of actions (encoded in vmPFC) may maintain, or even strengthen, optimistic bias. Yet, lowered integration of negative emotional memories may lead to emotional processing deficits (Baker et al., 2010; Townsend & Altshuler, 2012), potentially leading to the vicious cycle depicted in *Fig. 2&5*, and increase risk for mood dysregulation (Phillips et al., 2008; Sheppes et al., 2015), difficulties with close relationships (Kealy et al., 2011), and a tendency towards maladaptive behaviors, such as substance use (Berking et al., 2011; Le Berre, 2019), aggressivity, and hostility (Calvete & Orue, 2012; Cole & Zahn-Waxler, 1992). This compromised emotion processing resembles effects of trauma (Etkin & Wager, 2007) and chronic stress (Ragen et al., 2016).

**Figure 5.**
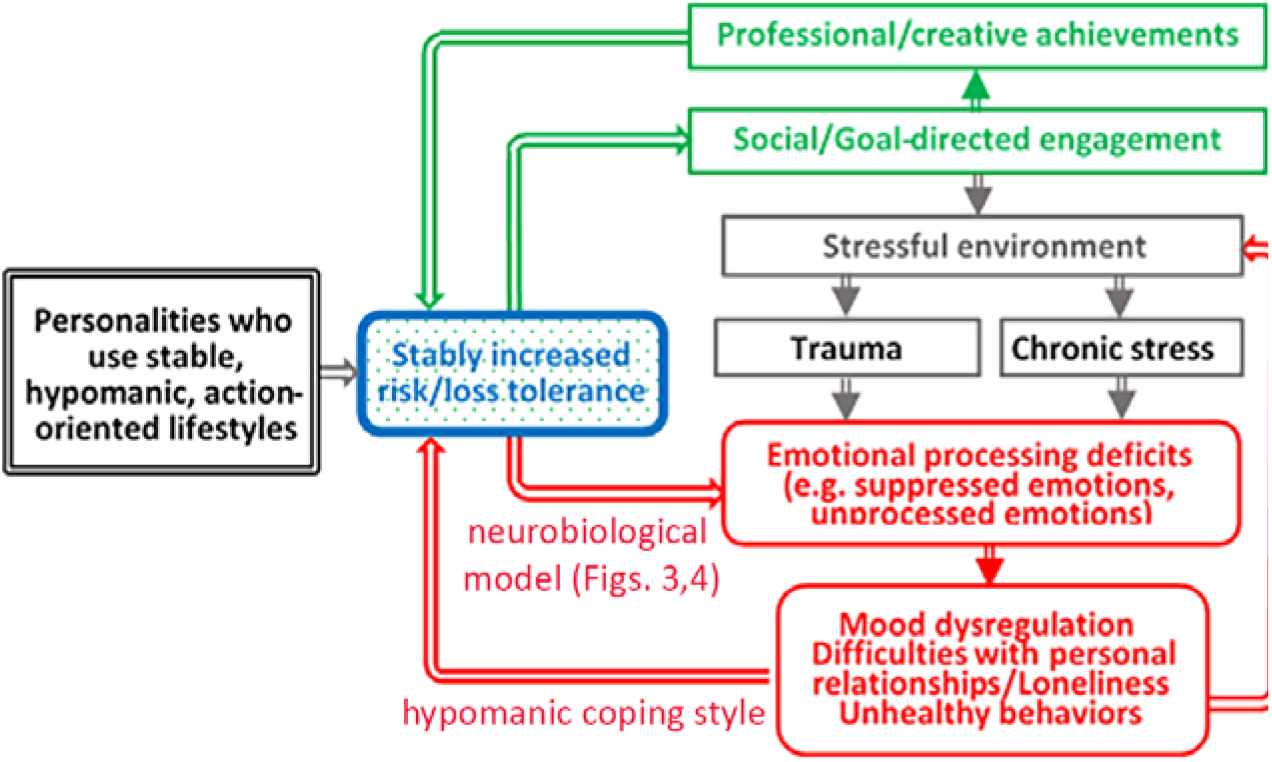
Dynamic model of interrelation between component of virtuous (**green**) and vicious (**red**) cycles in elevated hypomanic tendencies

Despite these costs, individuals with elevated hypomanic tendencies continue to engage in risky activities (Shein, 1974). Risk takings may expose these individuals to stress and trauma, further compromising emotional processing and feeding the vicious cycle (Harms, 2019). Individuals with elevated hypomanic tendencies show heightened AMY activation during passive viewing of negative pictures (Heissler et al., 2014) and are less successful in down-regulating AMY activity by reappraising negative stimuli (Heissler et al., 2014). This AMY dysregulation may be consistent with the possibility of progressively ‘overloaded’ limbic system (a term we borrow from the trauma literature (Dannlowski et al., 2012)) in hypomanic temperament (Stevens et al., 2018). Thus, results from our two studies suggest that increased risk-taking in the face of potential loss together with reduced integration of negative emotional experience may represent important candidate mechanisms underlying hypomanic vulnerability, potentially contributing over time to unprocessed negative affect and progressive cortico-limbic dysregulation (“limbic overload”).

Our novel hypothesis of hypomanic vulnerabilities also suggests strategies that may mitigate the risk of limbic system dysregulation to promote resilience. For instance, they might proactively engage in cognitive reappraisal (Troy & Mauss, 2011), which can improve affective, cognitive, and social outcomes (Cutuli, 2014). Cognitive reappraisal increases linearly with age during young adulthood (McRae et al., 2012); we predict that this developmental trajectory is altered in young adults with elevated hypomanic tendencies, who may ‘postpone’ attending to their negative emotional experiences.

Neurobiologically, cognitive-emotional integration (Meyer et al., 2019; Pessoa, 2010) and dispositional use of cognitive reappraisal (Picó-Pérez et al., 2018) have been linked to interaction (fc) of the anterior insula (AI) with limbic (AMY and Hipp) and cortical regions (including vmPFC and DLPFC). The use of cognitive reappraisal strategies may strengthen rAI connectivity with limbic and cortical areas (e.g. vmPFC), reducing subsequent limbic reactivity to salient stimuli (Cutuli, 2014). Active reappraisal has been shown to decrease self-reported emotional and AMY responses to negative stimuli (Denny et al., 2015). In Study 2, rAI↔vmPFC fc during rest I negatively correlated with rAMY sensitivity to potential losses during R&A task, further supporting this idea (*Fig. 6*). A recent study demonstrated that in PTSD, symptom improvements following therapy were associated with increased insula↔AMY fc (Fonzo et al., 2021); this is consistent with the idea that improved insula↔AMY fc may mitigate progressive cortico-limbic dysregulation (‘limbic overload’).

Dispositional use of cognitive-emotional integration is positively associated with the quality of interpersonal relationships (English et al., 2012; Fardis, 2005; Mazzuca et al., 2019; Shahar et al., 2019). Emotional regulation training has been shown to reduce interpersonal conflict in close relationships (Molajafar et al., 2015), possibly through the development of empathic concern (Laghi et al., 2018). We suggest that these relations are bidirectional; that is, quality personal relationships enhance the use of cognitive reappraisal strategies and provide implicit empathy training, thereby contributing to resilience in individuals with hypomanic tendencies. This would explain why individuals with both elevated hypomanic tendencies and poor social connectedness (Chapman et al., 1984) had greater rates of psychopathology and poor overall adjustment compared to those with elevated hypomanic tendencies with high quality social connectedness.

Importantly, our dynamic model of hypomanic vulnerabilities suggests qualitatively different strategies for altering illness trajectories than do unidimensional models of hypomanic vulnerabilities. From the latter perspective, risk mitigation would logically be targeted to reducing specific risk factors (neural, behavioral, or environmental). More complex models recognize that some risk factors may have corollary benefits (e.g., creativity and entrepreneurship in individuals with elevated hypomanic tendencies), and that altering them may have complex effects that reverberate through the nonlinear system. These models direct us instead to enhance resilience, to adaptively complement core personality characteristics, like elevated hypomanic tendencies, rather than seeking to suppress them.

## Conclusion

Together, these findings support a dynamic biosocial framework in which hypomanic vulnerability reflects not simply elevated reward sensitivity or impulsive approach behavior, but persistent goal-directed engagement coupled with reduced integration of negative emotional experience into subsequent behavioral regulation. Across longitudinal personality data and pilot neuroimaging findings, elevated hypomanic tendencies were associated with enduring coupling of partially opposing temperament dimensions together with distinct cortico-limbic dynamics during loss-related decision making. We propose that this combination of elevated loss tolerance and reduced emotional updating may, over time, contribute to progressive “limbic overload,” increasing vulnerability to emotional dysregulation and psychopathology while simultaneously supporting high engagement, creativity, and achievement. Importantly, the present framework generates experimentally testable predictions regarding the neural, behavioral, developmental, and social processes that may contribute both to vulnerability and resilience in individuals with elevated hypomanic tendencies.

## Limitations and Future Directions

Several limitations should be considered when interpreting the present findings. First, both studies relied on relatively modest sample sizes, particularly the pilot neuroimaging study, and the observed associations therefore require replication in larger and independent samples. Second, although the proposed cortico-limbic model demonstrated excellent fit within the current dataset, its generalizability across populations, developmental stages, and task contexts remains unknown and should be directly evaluated in future work. Third, the present findings are correlational and do not establish causal relationships among emotional processing, cortico-limbic dynamics, behavioral persistence under loss, and hypomanic vulnerability. Additional longitudinal, experimental, and computational studies will be necessary to determine whether the proposed mechanisms prospectively predict changes in emotional regulation, resilience, or progression toward psychopathology. More broadly, the Limbic Overload Hypothesis should be viewed as an initial dynamic framework intended to generate empirically testable predictions that can be refined, modified, or expanded as additional behavioral, neurobiological, and longitudinal evidence accumulates.

